# Weighted Kernels Improve Multi-Environment Genomic Prediction

**DOI:** 10.1101/2022.04.10.487783

**Authors:** Xiaowei Hu, Brett F. Carver, Yousry A. El-Kassaby, Lan Zhu, Charles Chen

## Abstract

Crucial to variety improvement programs is the reliable and accurate prediction of genotype’s performance across environments. However, due to the impactful presence of genotype by environment (G×E) interaction that dictates how changes in expression and function of genes influence target traits, prediction performance of genomic selection (GS) using single-environment models often falls short. Furthermore, despite the successes of genome-wide association studies (GWAS), the genetic insights derived from genome-to-phenome mapping have not yet been incorporated in predictive analytics, making GS models that use Gaussian kernel primarily an estimator of genomic similarity, instead of the underlying genetics characteristics of the populations. Here, we developed a GS framework that, in addition to capturing the overall genomic relationship, can capitalize on the signal of genetic associations of the phenotypic variation as well as the genetic characteristics of the populations. The capacity of predicting the performance of populations across environments was demonstrated by an overall gain in predictability up to 31% for the winter wheat DH population. Compared to Gaussian kernels, we showed that our multienvironment weighted kernels could better leverage the significance of genetic associations and yielded a marked improvement of 4-33% in prediction accuracy for half-sib families. Furthermore, the flexibility incorporated in our Bayesian implementation provides the generalizable capacity required for predicting multiple highly genetic heterogeneous populations across environments, allowing reliable GS for genetic improvement programs that have no access to genetically uniform material.

## 1. Introduction

Variety improvement programs are tasked to capturing heritable genomic response to selection across multiple growing environments and field seasons. While climatic uncertainty is outpacing variety development, the condition of global food, fuel and fiber insecurity has become more vulnerable (Feynman and Ruzmainkin 2007). In the face of diverse abiotic stresses, when considering genomic selection for variety improvement (GS, Meuwissen et al. 2001), reliable prediction of genotype performance across environmental variabilities has become increasingly critical.

However, selection using single-environment (SE) models becomes unreliable in the presence of genotype by environment (G×E) interaction (Burgueño et al. 2012; Crossa et al. 2017), due to the heterogeneity of genetic variance across environments, or imperfect genetic correlation of the same traits across sites/seasons (Crossa et al. 2004). Recently, GS models capable of assessing single population performance across multiple environments (ME) have been proposed (López-Cruz et al. 2015; Crossa et al. 2016; Lado et al. 2016; Montesinos-López et al. 2016; Spindel et al. 2016; Cuevas et al. 2018). In their study examples using two maize and two wheat data sets, Cuevas et al. (2018) examined the prediction accuracy of six different GS models with G×E interactions. Though some degree of advantage over the conventional SE counterparts can be identified, such gain can only be observed when the phenotypic correlation between environments was high (i.e., above 0.6), and when the traits of interest had moderate to high heritability (Burgueño et al. 2012; López-Cruz et al. 2015; Cuevas et al. 2016; Monteverde et al. 2018). The negative impact of the G× E to the performance of GS models is evident, even more so when inbred lines or homogeneous growing conditions are unavailable (Resende et al. 2012). For example, in tree genetic improvement programs that primarily use open-pollinated families, the half-sib pedigree structure has eventually prevented partitioning G×E interaction from genetic variance owing to the lack of clonal replications or genetically uniform lines, thus impeding the performance of GS models (Beaulieu et al. 2014; Chen et al. 2018; Gamel El-Dien et al. 2018; Alves et al. 2020; Thistlethwaite et al. 2020). To address this challenge in GS, here we developed a statistical model capable of assessing the prediction performance of multiple genetic populations across different environments. To our knowledge, this is also the first GS framework capable of predicting the performance of genetically heterogeneous populations across multiple environments.

Genome-wide approaches have played an instrumental role in the discovery of new biological insights underpinning complex trait variation (Frazer et al. 2009; Huang et al. 2010; Wang et al. 2019). Despite the tens of thousands of variant-trait associations cataloged (Buniello et al. 2019), knowledge learned from genome-wide association studies (GWAS) has not been adequately modeled in GS framework. Predicted by population genetics models, studies attempting to understand the impact of rare or less common variants on complex traits have shown an inverse relationship between the variant’s effect size and its frequency in the population (Park et al. 2011; Bomba et al. 2017). Empirical results from recent GWAS studies are mostly in agreement-that is, common variants have small effects, and rare variants have large effects (Bloom et al. 2019; Fournier et al. 2019; Wainschtein et al. 2019). Low-frequency and rare variants with small to modest effects that are thought to contribute to the missing heritability of many complex traits (Manolio et al. 2009; Eichler et al. 2010) may often have been overlooked because of the process in array production (Ziegler et al. 2008. Bouwman et al. 2017; Zhang et al 2018). As a consequence of not able to capture these rare but favorable alleles, selection based on the genomic estimated breeding values (GEBVs) could lead to loss of genetic diversity which further reduces the long-term genetic gain and prediction accuracy (Jannink 2010; Eynard et al. 2015; Liu et al. 2015; Doublet et al. 2019; Meuwissen et al. 2020; Vanavemaete et al. 2020).In this study, we proposed a flexible GS framework that incorporates marker information beyond just genotypic values, while extending the capability of conventional ME models. Our study uses examples in winter wheat and Interior spruce populations to demonstrate the advantage of including trait-and population-specific genetic characteristics, such as single nucleotide polymorphism (SNP) allele frequency and strength of association with the target phenotypes. Comparing to the existing Gaussian Kernel (GK) that assigns a uniform weight to every SNP, our proposed Weighted Kernel (WK) captured more robust genetic relationship of individuals within and cross environments by differentiating the contribution of SNPs. This capacity to address the trait- and population-specific environmental effects is not limited to the use of clonal genetic material or inbred lines, making modeling G× E in GS feasible for trials that utilize highly heterogeneous genetic resource to examine genetic adaptability across the range of a species or environmental variabilities (Risk et al. 2021).

High-throughput technologies have revolutionized biological and medical research and will continue to explore other omics space responsible for trait variation and adaptive responses to environmental variability (Halstead, et al. 2021; Hasin et al. 2017; Li, et al. 2019; Kim et al. 2016; Westhues et al. 2017). Integrating various omics information has become increasingly crucial for complex trait prediction and disease diagnostics (Tieri et al. 2011; Gomez-Cabrero et al. 2014; Higdon et al. 2015; Huang et al. 2017). The ability of kernelbased approaches to leverage the complementing favorable properties of predictors (Schrag et al. 2018), and the association significance to trait variation across growing conditions, would have the potential to provide consistent predictability across traits and environments.

## 2. Materials and Methods

### 2.1. Duster x Billings hard red winter wheat Doubled Haploid (DH) population

Developed cooperatively by the Oklahoma Agriculture Experiment Station (OAES) and the USDA-ARS, a total of 242 DH lines derived from the intercross of Duster and Billings winter wheat varieties were used in the study. Traits analyzed include grain yield (GY) in kilograms per hectare (kg/ha), sodium dodecyl sulfate sedimentation value (Lorenzo and Kronstad, 1987) adjusted for flour protein content (SDS), kernel weight measured by the single kernel characterization system (Perten Instruments, Segeltorp, Sweden) (SKCSKW), and wheat protein on a 12% moisture basis (WHTPRO). Each of these traits was evaluated in three harvest years with varied rainfall (*i.e.,* 19.8, 41.3, and 45.2 cm for 2014, 2015, and 2016, respectively) in Stillwater, OK, USA (36.12N, 97.09W), representing three different environments. The average of two field replicates of each DH line per year was used in the analysis.

Genotypes were derived using genotype-by-sequencing (GBS) technology, and 16,265 SNP markers were selected after filtering markers with >50% missing ratio. Missing genotypes of markers were imputed by the marker mean (Nazzicari et al. 2016). Although the genetic profile is the same across three years, the effects of environment on phenotype can vary in different years. Hence, we estimated the single-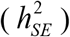 and multi-environment narrow-sense heritability 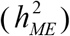 from GBLUP using single-year phenotypes and the average of three-year phenotypes for each trait, respectively. The estimation was implemented in the R package BGLR (Pérez and de los Campos, 2014).

### 2.2. Interior spruce population

The Interior spruce breeding population includes a total of 1,126 38-year-old trees growing over three sites in British Columbia Canada, *i.e.,* Prince George Tree Improvement Station (PGTIS), Aleza Lake, and Quesnel. Each site has 25 families with various sample sizes (Gamal El-Dien et al. 2015). To reduce the impact of unbalanced sample size on modeling, we randomized each family with respect to its minimum sample size (range from 6 to 16), resulting to 340 trees per site. Phenotypes used for prediction are height in m (HT) and diameter at breast height in cm (DBH) as growth traits; and two wood quality attributes, resistance to drilling (WD_res_) and wood density in kg/m^3^ using X-ray densitometry (WD_X-ray_). The genotypic information regarding GBS SNP can also be found in Gamal El-Dien et al. (2015). The single- 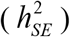 and multienvironment narrow-sense heritability 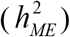 of each trait were estimated from GBLUP by Gamal El-Dien et al. (2015) and were reported in Table 1.

**Table 1.** Observed phenotypic correlation, single and multi-environment heritability estimates from genomic best linear unbiased prediction (GBLUP) for Duster x Billings winter wheat and the Interior spruce populations.

| | Trait | | E1 | E2 | E3 | $h_{SE}^2$ | $h_{ME}^2$ | Trait | E1 | E2 | E3 | $h_{SE}^2$ | $h_{ME}^2$ |
| --- | --- | --- | --- | --- | --- | --- | --- | --- | --- | --- | --- | --- | --- |
| **Wheat** | **GY** | E1 | 1 | 0.45 | 0.27 | 0.63 |  | **SDS** | 1 | 0.52 | 0.52 | 0.45 |  |
|  |  | E2 |  | 1 | 0.45 | 0.62 | 0.66 |  |  | 1 | 0.72 | 0.33 | 0.46 |
|  |  | E3 |  |  | 1 | 0.60 |  |  |  |  | 1 | 0.46 |  |
|  | **SKCSKW** | E1 | 1 | 0.28 | 0.40 | 0.46 |  | **WHTPRO** | 1 | 0.13 | 0.37 | 0.40 |  |
|  |  | E2 |  | 1 | 0.48 | 0.33 | 0.33 |  |  | 1 | 0.17 | 0.36 | 0.34 |
|  |  | E3 |  |  | 1 | 0.34 |  |  |  |  | 1 | 0.28 |  |
| **Spruce\*** | **HT** | E1 | 1 | 0.05 | 0.13 | 0.50 |  | **DBH** | 1 | 0.01 | 0.03 | 0.37 |  |
|  |  | E2 |  | 1 | 0.19 | 0.32 | 0.20 |  |  | 1 | 0.08 | 0.26 | 0.07 |
|  |  | E3 |  |  | 1 | 0.56 |  |  |  |  | 1 | 0.53 |  |
|  | **WD<sub>res</sub>** | E1 | 1 | 0.09 | -0.05 | 0.49 |  | **WD<sub>X-ray</sub>** | 1 | 0.19 | 0.05 | 0.28 |  |
|  |  | E2 |  | 1 | 0.08 | 0.28 | 0.10 |  |  | 1 | 0.16 | 0.39 | 0.18 |
|  |  | E3 |  |  | 1 | 0.42 |  |  |  |  | 1 | 0.43 |  |

Abbreviations: E1/E2/E3=Year 2014/2015/2016 for wheat; E1/E2/E3=PGTIS/Aleza Lake/Quesnel for spruce; PGTIS, Prince George Tree Improvement Station; $h_{SE}^2$ , heritability estimated from single-environment GBLUP; $h_{ME}^2$ , heritability estimated from multi-environment GBLUP; GY, grain yield; SDS, SDS sedimentation value; SKCSKW, kernel weight; WHTPRO, wheat protein; HT, height; DBH, diameter at breast height; WD<sub>res</sub>, resistance to drilling; WD<sub>X-ray</sub>, wood density in kg/m<sup>3</sup> using X-ray densitometry.
\* Heritability estimates, see Gamal El-Dien et al. (2015).

### 2.3. Statistical models

#### 2.3.1. Siangle-environment (SE) model

The kernel matrix in GS models is normally used to represent the genetic correlation between individuals that can be derived from either pedigree information or molecular marker data. To account for cryptic relatedness in genetic background, the SE model implemented here was an extension of the model 1 in Cuevas et al. (2017), by adding a random background genetic effect, **b**_*j*_. The SE model is expressed as follows:

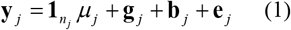

where **y**_*y*_ is the response vector with length *n_j_, n_j_*, is the total number of phenotypic observations in the *j^th^* environment, *j* = 1,!, *m, m* is the number of environments; **1***_n_j__* is a vector of ones with length *n_j_*, *μ_j_* is the overall phenotypic mean of individuals in the *j^th^* environment; **g**_*y*_ is the random genetic effect of individuals in the *j^th^* environment, and we assume 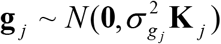 where 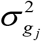 is the genetic variance of individuals in the *j^th^* environment, **K**_*j*_ (size *n_j_ × n_j_*) is the kernel matrix used to describe the genetic similarity between individuals in the *j^th^* environment; **b**_*j*_ is the random background genetic effect of the *j^th^* environment that is not explained by the genetic markers in the **g**_*j*_, and we assume 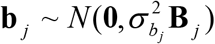 where 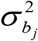 is the background genetic variance of individuals in the *j^th^* environment, **B**_*j*_ (size *n × n*.) is a matrix representing the background genetic relationship of two individuals in the *j^th^* environment; **e**_*j*_ is the random error term of the *j* environment, and we assume 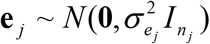 where 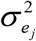 is the residual variance of the *j^th^* environment and *I_n_j__* is the identity matrix with size *n_j_*; **g**_*j*_, **b**_*j*_ and **e**_*j*_ are assumed to be independent.

#### 2.3.2. Multi-environment (ME) model

To fully capture G× E interaction, we proposed a generalization of the model 3 in Cuevas et al. (2017) as follows:-

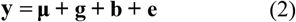

where 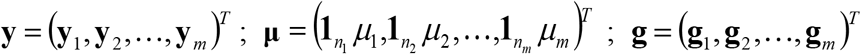 and 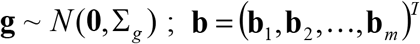 and 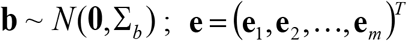 and **e** ~ *N* (**0**, Σ_*e*_); **g**, **b** and **e** are assumed to be independent;, **y**_*m*_, *μ_m_*, **g**_*m*_, **b**_*m*_ and **e**_*m*_ are defined the same as in SE model.

In general, the genetic covariance matrix is

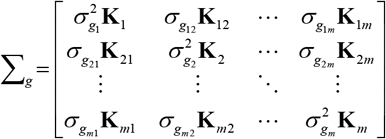

where 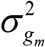 and **K**_*m*_ are defined the same as in SE model; 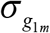 is the genetic covariance of individuals in the 1^*st*^ environment and *m^th^* environment; **K**_1*m*_ (size *n*_1_ × *n_m_*) is the kernel matrix representing the genetic relationship between individuals from the 1^*st*^ and *m^th^* environment.

The background genetic covariance matrix is

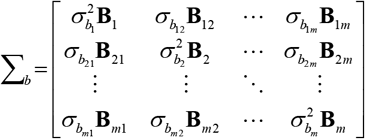

where 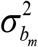 and **B**_*m*_ are defined the same as in SE model; 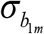 is the background genetic covariance of individuals in the 1^*st*^ and the *m^th^* environment; **B**_1*m*_(size *n*_1_ × *n_m_*) is a matrix constructed to present the background genetic relationship between two individuals in the 1^*st*^ and *m^th^* environment. With this model, background genetic relationship can be appropriately incorporated into both SE and ME models. For instance, when two individuals from the same family in the Interior spruce population, a half-sib relatedness of 0.25 was assigned to indicate their shared genetic background. While when the informative background relationship is unavailable, an identity matrix can be used for both background genetic variance and covariance matrices (Crossa et al. 2017), which was the case for Oklahoma wheat DH population demonstrated in this study.

The covariance matrix of the residuals is

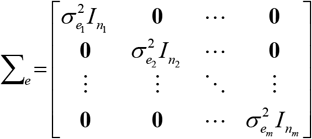

where 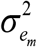 and *I_n_m__* are defined the same as in SE model.

#### 2.3.3. Gaussian kernel (GK) and Weighted kernel (WK)

Here in this study, we compared the model prediction performance with two different kernels, Gaussian kernel (GK) used in de los Campos et al. (2010) and our proposed weighted kernel (WK). The GK in (3) transforms genetic distance into genetic correlation between individuals.

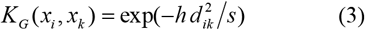

where *K_G_*(.,.) is a positive definite function evaluated by marker genotypes; *x_t_,x_k_* are vectors of marker genotypes for the *i^th^* and *k^th^* individuals respectively, *i,k* = 1,...,*n_j_*, *x_i_* =(*x*_*i*1_,...,*x*_*i*l_,..., *x_ip_*)^*T*^ and *x_k_* =(*x*_*k*1_,...,*x_kl_*,..., *x_kp_*)^*T*^, *l* = 1,...,*p*, *p* is the total number of markers; the allelic states of *x_il_* are coded as 0, 1, 2 for AA, Aa, and aa respectively; *h* is a positive bandwidth parameter that controls the rate of decay of the genetic correlation between two individuals. To determine the optimal value of the parameter, either a grid search method from cross-validation procedure or an empirical Bayesian approach (Pérez-Elizalde et al. 2015) can be applied. In this study, *h* = 1 was used for simplicity. 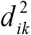 is squared Euclidean distance between two individuals *i* and *k* explained by marker genotypes, and *s* is the sample maximum of 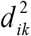.

Motivated by Wu et al. (2011) and Yan et al. (2014), we proposed to model additional information such as the frequency and the effects of the variants by a WK method in (4).

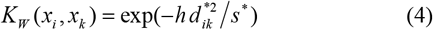

where 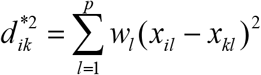, *s*\* is the sample maximum of 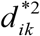, and the weight *W_l_* assigned to the *l^th^* marker is based on its minor allele frequency (MAF) and p-values from single-marker association (SMA) test of model with G×E interaction. The detailed formula of *w_l_* is the following

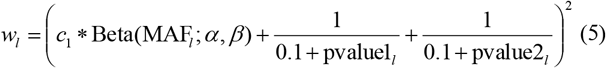

where *c*_1_ is a constant; MAF_*l*_ is the minor allele frequency of the *l^th^* marker; *α* and *β* are the parameters of Beta distribution density function; pvalue1_*l*_ and pvalue2_*l*_ are p-values of main genotypic effect and G×E interaction effect respectively from SMA test for the *l^th^* marker, these p-values are adjusted by false discovery rate at 0.05.

To account for the potential effects of low frequency variants while incorporating the test statistics from SMA test, we proposed the following formula (6) to determine the value of *c*_1_.

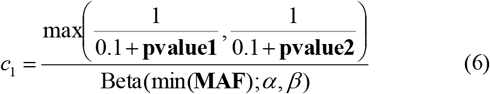

where **pvalue1**= (*pvalue*1_1_,...,*pvalue*1_*l*_,..., *pvalue*1_*p*_)^*T*^, **pvalue2**= (*pvalue*2_1_,..., *pvalue*2_*i*_,...,*pvalue2*_*p*_)^*T*^, and **MAF**= (*MAF*_1_,..., *MAF*_l_*MAF_P_*)^*T*^.

As for the setting of *α* and *β* in (5) and (6), Wu et al. (2011) and Yan et al. (2014) suggested to set *α* = 1 and *β* = 25 as a general way to control the impact from rare genetic variants in their GWAS research. In this study, we proposed to fix *α* = 1 and explore the impact of *β* on the performance of prediction as such beta density decreases as MAF increases (see Fig. S1 for details). As expected, when both MAF and p-value are very small, the value of 10 *c*_1_ was found to be determined approximately by *β*, *i.e*., 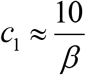. As a result, *β* cannot go to infinity to shrink *c*_1_ towards zero. To further document the impact of *β* on prediction performance of model with WK, five values were 10 inspected, *i.e., β* = 12, 25, 50, 100, and 200. Thus, 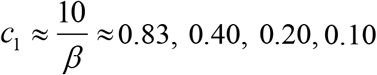, and 0.05.

In addition, we compared the prediction performance of the model using the proposed WK with the model that implements the WK by MAF or p-value alone, denoted as WK_MAF_ and WK_Pvalue_ respectively. As comparison, we denoted the WK contributed by both MAF and P-value as WK_MAF_Pvalue_, and its weight *w_l_* is from equation (5). The weight in WK_MAF_ is calculated by equation (7).

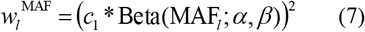

where *c*_1_,*α*, and *β* are defined the same as above.

Similarly, the weight in WK_Pvalue_ is formulated as following

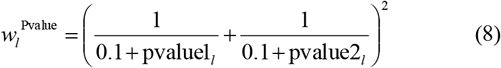

### 2.4. Model Implementation

#### 2.4.1. SE model

Analysis was conducted in R (R Core Team, 2020). The SE model was implemented using R package BGLR with 12,000 iterations and the first 6,000 as burn-in for both wheat and spruce data sets (Pérez and de los Campos, 2014).

#### 2.4.2. ME model

##### 2.4.2.1. Duster x Billings hard red winter wheat DH population

In the cases where clonal material is used in multiple environments, such as Oklahoma winter wheat DH breeding populations, the ME model was fitted using R package MTM (de los Campos and Grüneberg, 2016) with 20,000 iterations and the first 10,000 samples as burn-in.

##### 2.4.2.2. Interior spruce population

Since no clonal or inbred line material were available for Interior spruce, we expanded the ME model for such scenario by a Bayesian approach to estimate all parameters in the ME model. The detailed implementation of Bayesian approach for ME model can be seen in the **Supplementary Material**. The model was implemented in R with 100,000 iterations and the first 50,000 as burn-ins. The convergences of Markov Chains for all models were assessed by visualizing the trace plots and running convergence diagnosis using R package CODA (Plummer et al. 2006).

#### 2.4.3. Prediction accuracy evaluation

Both Pearson’s correlation coefficient (PCOR) between observed and predicted phenotypes, and its mean squared error (MSE) were used to assess the model prediction accuracy. To train the model, we first split the whole data into a training (TRN) and a testing (TST) data sets, we randomly assign 70% of the data as TRN and the remaining 30% as TST for the evaluation of SE models. For ME models, we applied cross-validation 2 (CV2) scheme that mimics the practical prediction scenario related to plant breeders where individual plants are only tested in some environments (Burgueño et al. 2012). The prediction of TST, as a consequence, will gain strength from both within- and cross-environment TRN that are related to TST. The procedure proposed in López-Cruz et al. (2015) was followed in this study CV2 scheme. Then the random partition for all models was repeated 50 times to generate an average prediction performance.

## 3. Results

For each dataset, we present the following: 1) MAF distribution; 2) summary of phenotypes and the estimated heritability; and 3) prediction performance of the proposed models in the single- and multi-environment settings. PCORs were used to illustrate the prediction performance. Additionally, the results of MSE were found to be consistent with PCORs, *i.e.,* a lower MSE tends to have a higher PCOR. The detailed evaluation of prediction performance can be found in the **Tables S1-S2**.

### 3.1. Minor allele frequency distribution

#### 3.1.1 Duster x Billings hard red winter wheat DH population

The distribution of common and rare SNP allele frequency for Duster x Billings DH population is shown in **Fig. 1A**, The wheat DH population has ~ 64% of the SNPs with MAF < 0.2, about 59% < 0.1 and 19% between 0.4 to 0.5. Further demonstrated in **Fig. S1**, the density of *Beta* (1; *β*) will converge to zero with large value of Beta random variables. Additionally, regarding to the context of MAF as the value of Beta random variable, 0.2 was considered as the large value.

**Figure 1.**
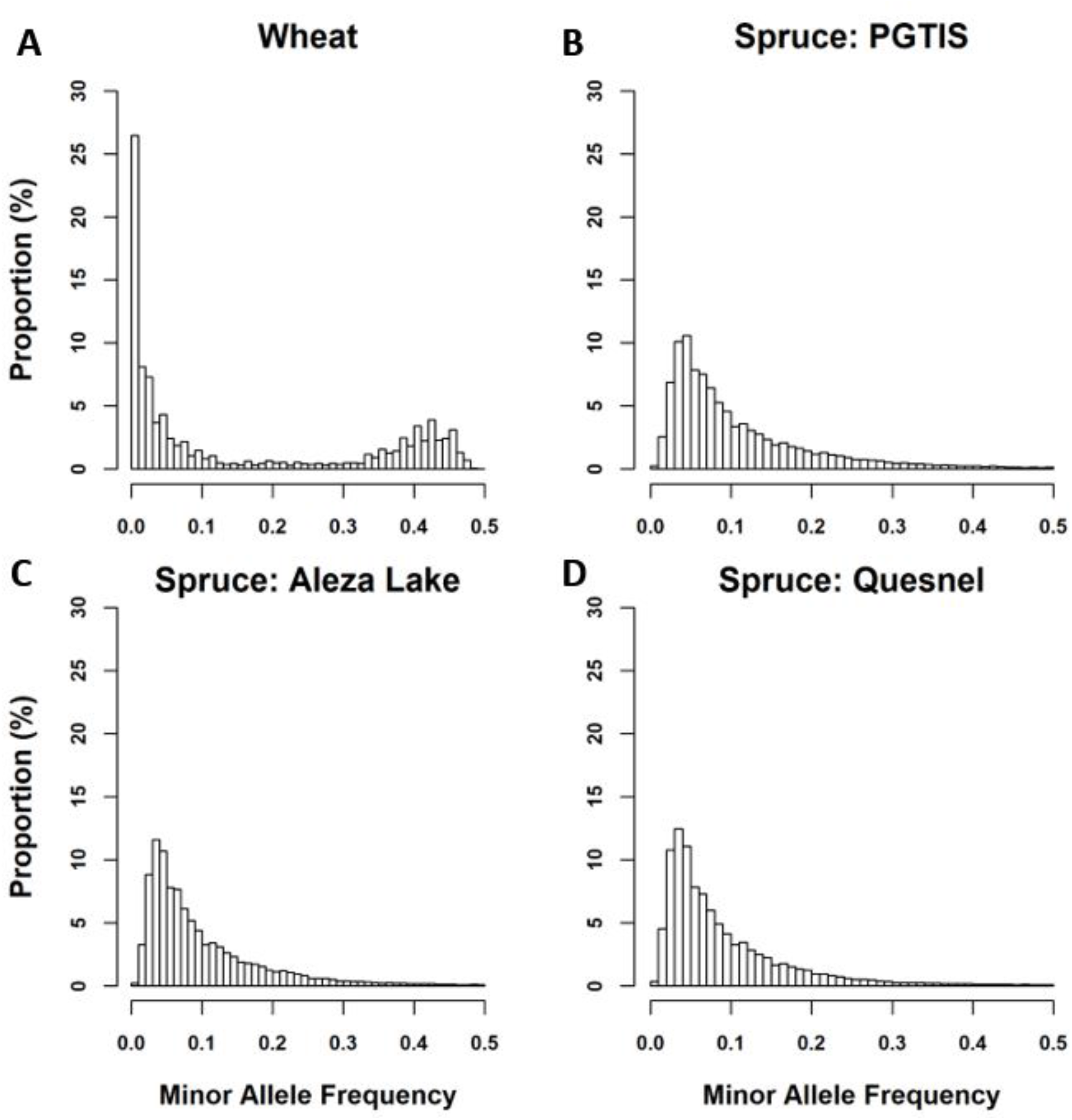
The distributions of minor allele frequency from genomic data of red hard winter wheat doubled haploid (DH) and Interior spruce populations. PGTIS, Prince George Tree Improvement Station.

#### 3.1.2. Interior spruce population

The distribution of MAF was found with a greater degree of rare alleles in all three sites of Interior spruce. There were about 86, 88, and 91% of SNPs with MAF < 0.2, for, PGTIS, Aleza Lake, and Quesnel, respectively (**Fig. 1B-D**).

### 3.2. Summary of phenotypes and heritability estimates

#### 3.2.1 Duster x Billings hard red winter wheat DH population

Boxplots of the four traits (GY, SDS, SKCSKW, and WHTPRO) across the three years (2014-2016) for Oklahoma winter wheat are shown in **Fig. 2A**. Both trait distributions and phenotypic variation are quite different among the three years for all four traits. For observed phenotypic correlation between years, SDS exhibited the highest average phenotypic correlation (0.52-0.72, average = 0.59), while WHTPRO was the lowest (0.13-0.37, average = 0.22) (**Table 1**). Heritability for each year 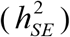, as well as cross-year estimates 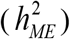, are also listed in **Table 1**. GY showed the highest and the most stable single-year heritability among the studied four traits 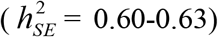. Conversely, the other three traits had much lower and varying heritability estimates. For multiyear heritability estimates (2014-2016), only GY and SDS have higher heritability than their single-year estimates (*i.e*., 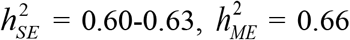 for GY; 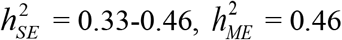 for SDS).

**Figure 2.**
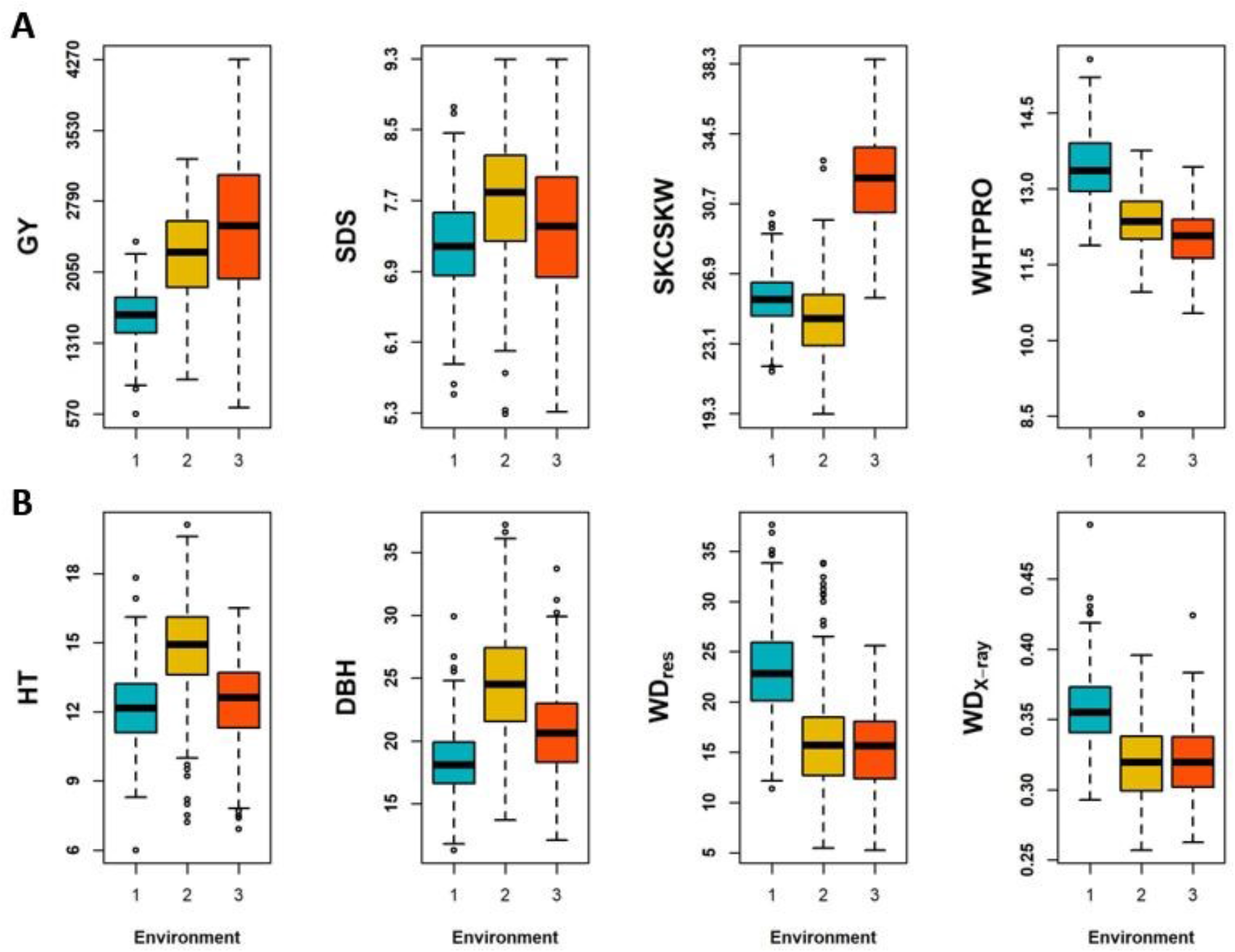
Boxplots of phenotypes. **A**) Winter wheat, grain yield (GY), SDS sedimentation value (SDS), kernel weight (SKCSKW), and wheat protein (WHTPRO) in each environment (Environment 1/2/3 = Year 2014/2015/2016); **B**) Interior spruce, height (HT), diameter at breast height (DBH), resistance to drilling (WD_res_), and wood density in kg/m^3^ using X-ray densitometry (WD_X-ray_) in each environment (Environment 1/2/3 = PGITS/Aleza Lake/Quesnel, PGTIS, Prince George Tree Improvement Station).

#### 3.2.2. Interior spruce population

Growth phenotypes, HT and DBH, varied among the three Interior spruce sites, while traits related to wood density *(e.g.,* WD_res_ and WD_X-ray_), Aleza Lake and Quesnel showed similarity in distribution and ranges (**Fig. 2B**). However, unlike the winter wheat population, the observed pairwise phenotypic correlations were relatively low for all studied traits (**Table 1**). Overall, WDX-ray had the highest average phenotypic correlation at 13% over the three sites (5-19%), and the lowest was found in the correlation with DBH (1-8%, average at 4%). The single-site heritability ranged from moderate to high (*i.e*., 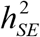 ranged from 0.26-0.56). Generally, traits measured in Quesnel showed higher heritability than the other two sites. The overall heritability estimated across the three sites were reduced to 0.07-0.20 (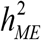, **Table 1**), with the highest in HT and lowest in DBH.

### 3.3. Single and multi-environment predictions

For the performance of WK, we present the results with *β* that produced the highest prediction accuracy of using WK_MAF_Pvalue_ for both data, *i.e., β* = 12for wheat (**Fig. S2**) and *β* = 200 for spruce (**Fig. S3**).

#### 3.3.1. Duster x Billings hard red winter wheat DHpopulation

The average prediction accuracies of SE and ME models are shown in **Fig. 3** for the DH hard red winter wheat population. In general, significant improvement of prediction accuracy can be seen with modeling across multiple environments (**Fig. 3**). For example, the prediction accuracy of GY using single year Gaussian kernel (SE_GK, in **Fig. 3**) ranged from 0.38 (2016) to 0.55 (2014). With the same Gaussian kernel, ME_GK trained the model with data from all years and generated 6-10% improvement in GY prediction accuracy. The gain from ME models can be as significant as a four-time increase (0.1 in SE_GK and 0.38 for ME_GK, for the SKCSKW in **Fig. 3**); substantial increase in ME_GK prediction accuracy was also found for SDS with an average increase of 35% over the SE_GK (**Fig. 3**). SDS also showed the highest gain of the estimated genetic variance from ME_GK vs. SE_GK (**Table S3**).

**Figure 3.**
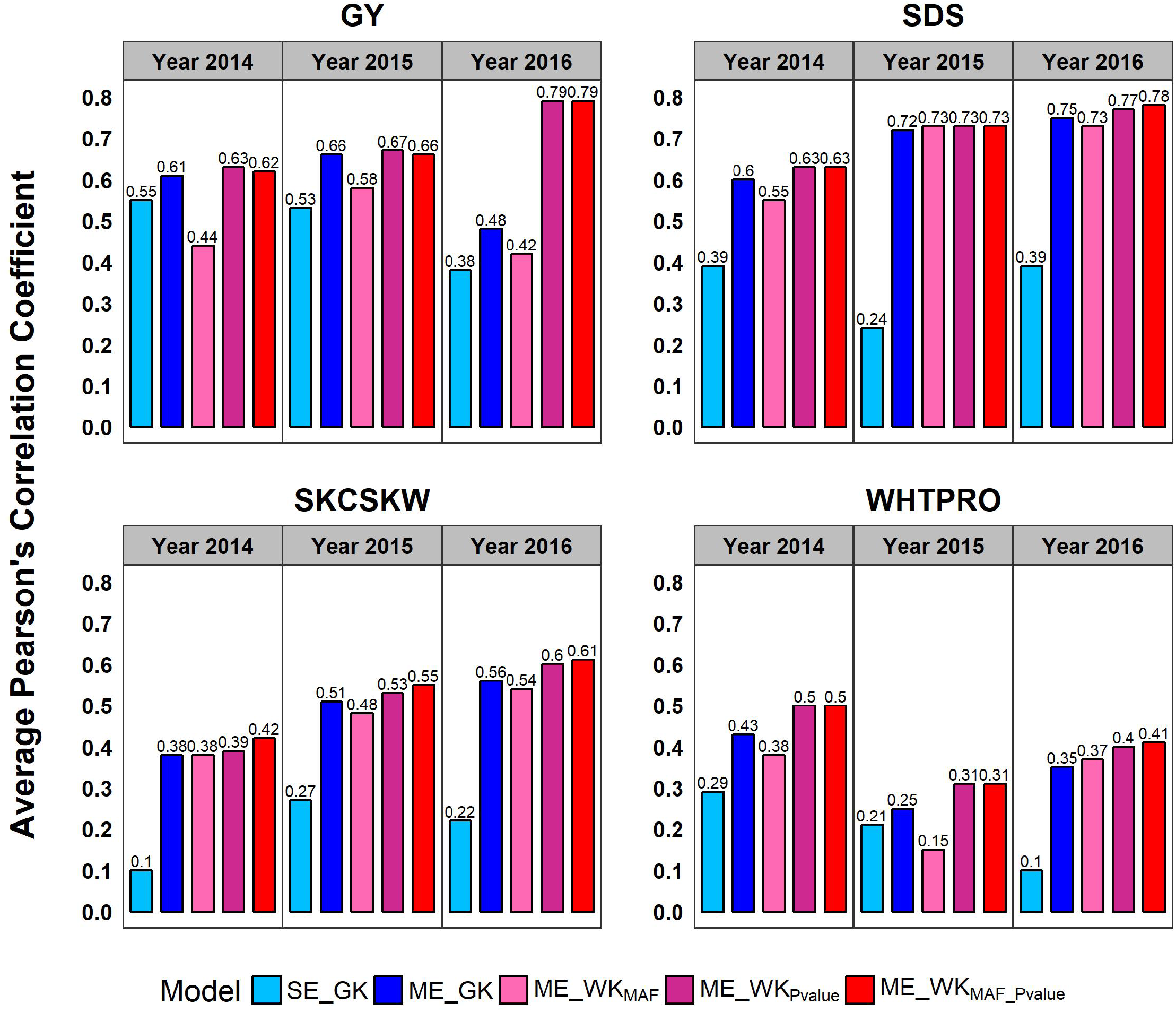
Prediction performance of genomic selection models for winter wheat. Average Pearson’s correlation coefficients were collected over 50 replications of CV2 scheme; SE_GK, single-environment model with Gaussian kernel; ME_GK, multi-environment model with Gaussian kernel; ME_WKMAF, multi-environment model with weighted kernel (WK) by minor allele frequency (MAF); ME_WK_Pvalue_, multi-environment model with WK by genome-wide association study p-value; ME_WK_MAF_Pvalue_ (with *β* = 12), multi-environment model with WK by both MAF and p-value; GY, grain yield; SDS, SDS sedimentation value;, SKCSKW, kernel weight; WHTPRO, wheat protein.

Weighting with MAF and the association signal further improved the prediction performance by WK for ME models, with the exceptions of the reduced accuracy found in the ME_WK_MAF_ model for GY and WHTPRO (**Fig. 3**). Among all three methods of WK, WK_MAF_Pvalue_ performed similarly to WK_Pvalue_, and both significantly outperformed WK_MAF_ for all studied traits. Compared with the ME_GK models, the increase of prediction accuracy from ME_WK_MAF_Pvalue_ models ranged from 1 to 3%, 4 to 5%, and 6 to 7% for SDS, SKCSKW, and WHTPRO, respectively (**Fig. 3**). For GY, the performance of ME_WK_MAF_Pvalue_ and ME_GK was found similar in 2014 and 2015, but ME_WK_MAF_Pvalue_ produced significantly higher prediction accuracy for 2016 (*i.e,* 31% higher than ME_GK, **Fig. 3**).

#### 3.3.2. Interior spruce population

Different from what observed in the wheat dataset, the advantage of ME_GK over SE_GK was not as evident for Interior spruce, which might be a result of the large amount of variance that cannot be accounted for in the multienvironment models (**Table S4**). The highest prediction accuracy for SE_GK was found in Quesnel for HT (**Fig. 4**); similar performance in HT was found for ME_GK as well. Addiitionally, wood quality traits showed consistent prediction accuracy for all Gaussian models, ranging from 0.19 to 0.31 for WD_res_ and 0.24 to 0.28 for WD_X-ray_. In **Table 1**, DBH had the lowest multi-environment heritability estimates, which is further reflected on the average of 5% reduction in ME_GK prediction performance (**Fig. 4**).

**Figure 4.**
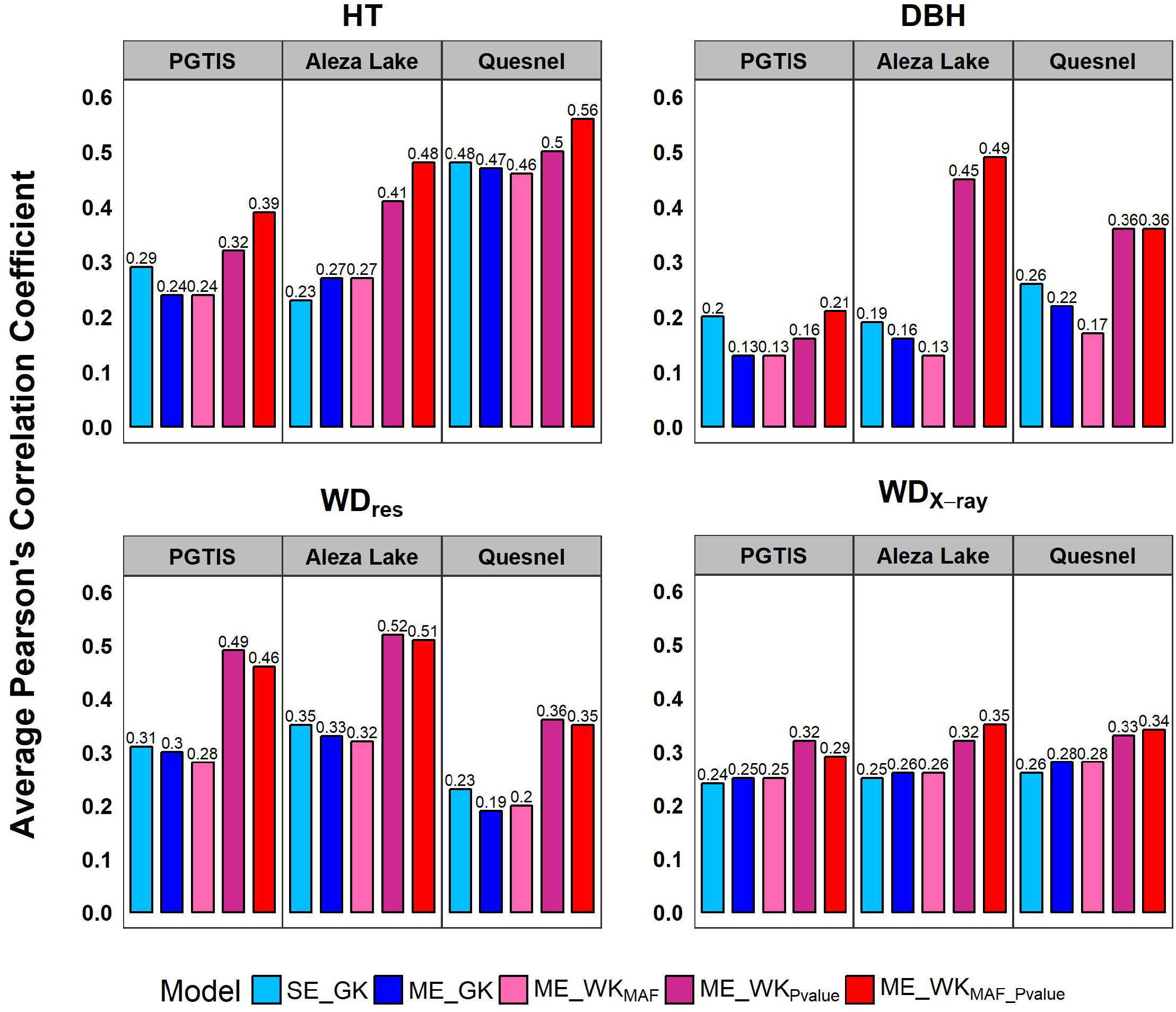
Prediction performance of genomic selection models for the Interior spruce population. Average Pearson’s correlation coefficients were collected over 50 replications of CV2 scheme; SE_GK, single-environment model with Gaussian kernel; ME_GK, multi-environment model with Gaussian kernel; ME_WK_MAF_, multi-environment model with weighted kernel (WK) by minor allele frequency (MAF); ME_WK_Pvalue_, multi-environment model with WK by genome-wide association study p-value; ME_WK_MAF_Pvalue_ (with *β* = 200), multi-environment model with WK by both MAF and p- value; HT, height; DBH, diameter at breast height; WD_res_, resistance to drilling; WD_X-ray_, wood density in kg/m^3^ using X-ray densitometry; PGTIS, Prince George Tree Improvement Station.

In general, modeling with MAF and specific-trait association improved predictability, even when predicting phenotypes for genetically heterogeneous material across environments. Among the WK implementations, WK_MAF_Pvalue_ outperformed the other WK models almost in all traits, and the WK_Pvalue_ model showed slight advantage for WD_res_ prediction in the PGTIS site (**Fig. 4**). HT was the most predictable phenotype, with a moderate prediction accuracy in Quesnel, increased from 0.48 of SE_GK and 0.47 of ME_GK to 0.56 ME_WK_MAF_Pvalue_. The greatest gain by using the WK models was, however, found in DBH in Aleza Lake; the benefit of using WK models for DBH was, however, diminished in PGTIS (**Fig 4**). The benefit of including genomics signals was not significant for WD_X-ray_. Due to the relatively indifferent prediction performance for WD_X-ray_ across all models, the benefit of incorporating MAF and association signal was not observed. We suspect that the SNP predictors generated for Interior spruce are in weak LD with the underlying genes and QTLs.

## 4. Discussion

GS performance can be influenced by many interrelated factors, including trait genetic architecture, heritability, and the relatedness among individuals in the populations (Crossa et al. 2017). When ME prediction across sites or growing seasons was conducted with a more defined set of genetic diversity like populations derived from controlled crosses, the advantage of incorporating available genetic correlation between environments was evident. As shown in **Fig. 3**, our ME_GK models using conventional GK demonstrated a consistent improvement over the SE model, showing a 4-38% gain in predictability for Oklahoma winter wheat DH population. The greatest improvement for this population was observed in SDS for 2015, the trait that also showed the most consistent cross-year prediction in Hu et al. (2019). Our results demonstrated that, even in the presence of identifiable environmental variability (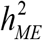 ranges from 0.33 to 0.66, **Table 1** and **Table S3**), the benefit of employing ME prediction can be anticipated in this case, because of the model capacity to leverage genotype’s environment-specific effect.

Shown in **Fig. 4**, the ME_GK model, on the other hand, exhibited a slightly unfavorable performance for Interior spruce, except for WDX-ray whose accuracies were found indifferent with the SE model. Compared to our results, the prediction analysis using the same half-sib families in Gamel El-Dien et al. (2015) presented a much-reduced GS accuracy with cross-site validations, even when the prediction accuracy was calculated by correlating the breeding values with the GEBVs. The non-additive effect of these traits was found significant in Gamel El-Dien et al. (2018), with WD_X-ray_ being the only exception. In this study, the multi-site heritability estimates 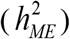 ranged from 0.07 to 0.20 (**Table 1**); this small amount of additive genetic variance would be one of the leading attributes that hinders the performance of the ME_GK model.

The conventional GK using genetic markers is only able to capture the overall genetic similarity between individuals. Although the bandwidth parameter in GK can adjust the distribution of genetic similarity (Pérez-Elizalde et al. 2015), such tuning is uniform to all genetic markers. For Interior spruce, the genetic marker data revealed a much lower relatedness of these trees within each site, suggesting the actual sibling relatedness within families rarely met the half-sib relatedness assumption. In the case where various degree of genetic relatedness between individuals exists within the same family across sites, the strength of incorporating genetic correlation in the ME models using only the Gaussian kernel might be confounded by the heterogeneous genetic background, resulting in an accuracy slightly lower to the SE models (**Fig. 4**).

The bandwidth parameter tuning in conventional kernel models could potentially create a better mapping between the overall genetic distance among individuals to the phenotypic variation (Pérez-Elizalde et al. 2015). However, it does not reflect the trait’s genomic functional space, leaving important biological insights, such as allele frequencies and the underlying genetic architecture, out of the genome-to-phenome mapping in the GK models. GWAS studies have been a powerful tool to assessing the association between genetic variants and trait variations. The genetic variants identified indicate their functional roles or a close linkage with important genetic determinants for the traits of interest (Wu et al. 2011; Yan et al. 2014; Lin et al. 2016). Several studies have suggested prioritizing GWAS variants when creating the genomic relationship matrix could improve SE predictability of unrelated individuals (de los Campos et al. 2013; Ober et al. 2015; Morgante et al. 2018).

Despite the increases in GWAS statistical power afforded in large international consortia (Willer et al. 2013; Wood et al. 2014; Liu et al. 2015; Astle et al. 2016; Bomda et al. 2017), GWAS still only accounts for a fraction of heritability for most complex traits, a well-known phenomenon called “missing heritability” (Manolio et al. 2009). Genetic variants outside of the reach of the GWAS statistical power are considered to also contribute to the missing heritability (Speed et al. 2012), including common variants with weak effects, low-frequency (MAF 1-5%), and rare variants (MAF <1%) of small to modest effects, or their combination (Agarwala et al. 2013). When the true causative genetic variants remain unknown, GS has been proven more effective than classic marker-assisted selection. This is because GS employs all available markers as a ‘compete modeling’ methodology for estimating trait performance (Jia 2017). Compared to phenotypic selection, GS could lead to the acceleration of annual inbreeding rate and the loss of genetic diversity as it encourages selecting individuals with high GEBV early in variety improvement programs and those closely related to the training populations (Bassi et al. 2016; Doekes et al. 2018; Forutan et al. 2018). In order to provide stable predictability across populations, GS might also contribute to rapid fixation of genomic regions where consistent marker effects across populations can be identified (Clark et al. 2011; Pszczola et al. 2012; Allier et al. 2019). When the breeding decision is made to optimize short-term genetic gain with conventional GS, rare but favorable alleles could be overlooked. That will essentially reduce selection accuracy and genetic gain in the long term. Demonstrated in simulation studies, up-weighting such alleles would provide 8 to 30.8% greater long-term gain than that of un-weighted prediction methods (Jannink 2010; Liu et al. 2015), further advocating the WK approaches proposed in this study for the long-term reliability of GS (Rutkoski et al. 2015; Zhang et al. 2018; Ramasubramanian and Beavis 2021).

Here, we presented a flexible GS framework capable of incorporating important genetic attributes to breeding populations and trait variability while addressing the shortcomings of conventional GS models. Shown in **Fig. 3–4**, the advantage of incorporating trait- and population-specific genetic characteristics, like *p*-values of GWAS and MAF, was evident. The MAF component in our WK models aided in preserving the rare favorable variants, which are usually underpowered in GWAS. In addition, the WK considers the contribution of genetic markers to the trait-specific G×E. By further differentiating the effects of SNPs between growing environments, GS predictability can be improved for all traits studied for DH genotypes, as well as for the half-sib families of Interior spruce with considerable degree of environmental variability across sties. Finally, the Bayesian kernel methodology presented in the present study offers the flexibility required for predicting multiple populations across environments without using genetically clonal material. This implementation, to our knowledge, is the first GS framework capable of predicting the performance of highly genetically heterogeneous populations across environments.

## Acknowledgements

Funding for this work was supported by grants from the Oklahoma Wheat Research Foundation (for XH, BFC and CC), Oklahoma Center for the Advancement of Science and Technology (OCAST) award number PS15-011-2 and PS19-004 for CC. This research represents the research outcomes for the USDA HATCH project OKL03011 (CC). Genotyping effort of this manuscript was also supported from the National Science Foundation award number NSF-MRI 1626257 (CC). This work was completed utilizing the High-Performance Computing Center facilities of Oklahoma State University at Stillwater, and also in part by the Extreme Science and Engineering Discovery Environment (XSEDE), which is supported by National Science Foundation grant number ACI-1548562. Specifically, it used the Bridges system, which is supported by NSF award number ACI-1445606, at the Pittsburgh Supercomputing Center (PSC) under the resource allocation MCB-180177.

## Data and Code Availability

Code for our proposed model are available here https://github.com/XiaoweiHu-Stat/Multivariate_WeightedKernel

## Supplementary Material

Supplementary Table S1. Average mean squared error (standard deviation in parentheses) of genomic selection models for winter wheat DH population.

Supplementary Table S2. Average mean squared error (standard deviation in parentheses) of genomic selection models for Interior spruce.

Supplementary Table S3. The estimated variance component proportions from genomic selection models for winter wheat DH population.

Supplementary Table S4. The estimated variance component proportions from genomic selection models for Interior spruce.

Supplementary Figure S1. The relationship between minor allele frequency and the weight from minor allele frequency by different values of *β* for winter wheat DH population and site PGTIS of Interior spruce.

Supplementary Figure S2. Average Pearson’s correlation coefficients over 50 replications of CV2 scheme from multi-environment model with weighted kernel by both minor allele frequency and p-value (ME_WKMAF_Pvalue) and from different values of *β* for winter wheat DH population.

Supplementary Figure S3. Average Pearson’s correlation coefficients over 50 replications of CV2 scheme from multi-environment model with weighted kernel by both minor allele frequency and p-value (ME_WKMAF_Pvalue) and from different values of *β* for Interior spruce.

