## Supplementary material for "Weighted Kernels Improve Multi-Environment Genomic Prediction": MultiEnvWK_SuppMaterial_XH.pdf

**Supplementary Material for Association Genomics Knowledge Guided Multi-Environment Genomic Prediction**

### Methods

#### Multi-environment (ME) model algorithm for heterogeneous populations

For heterogeneous populations scenario, we apply Bayesian method to estimate all parameters in ME model. In general, $\mathbf{y} \mid \mu, \mathbf{g}, \mathbf{b}, \mathbf{e} \sim N(\mu, \Sigma)$ where $\Sigma = \Sigma_g + \Sigma_b + \Sigma_e$. And the estimated parameters are $\theta = (\mu_1, ..., \mu_m, \sigma^2_{g_1}, ..., \sigma^2_{g_m}, \sigma_{g_{12}}, ...\sigma_{g_{jm}}, ..., \sigma^2_{b_1}, ..., \sigma^2_{b_m}, \sigma_{b_{12}}, ...\sigma_{b_{jm}}, ..., \sigma^2_{e_1}, ..., \sigma^2_{e_m})^T$. However, we reparametrize $\sigma_{g_{jm}} = \rho_{g_{jm}} \sigma_{g_j} \sigma_{g_m}$, then the actual parameters estimated in the model are $\theta = (\mu_1, ..., \mu_m, \sigma^2_{g_1}, ..., \sigma^2_{g_m}, \rho_{g_{12}}, ...\rho_{g_{jm}}, ..., \sigma^2_{b_1}, ..., \sigma^2_{b_m}, \rho_{b_{12}}, ...\rho_{b_{jm}}, ..., \sigma^2_{e_1}, ..., \sigma^2_{e_m})^T$. The joint posterior density of the parameters becomes:

$$\begin{aligned} P(\theta \mid \mathbf{y}) &\propto P(\mathbf{y} \mid \theta) P(\theta) \\ &\propto P(\mathbf{y} \mid \theta) P(\mu_1) \cdots P(\mu_m) P(\sigma^2_{g_1}) \cdots P(\sigma^2_{g_m}) P(\rho_{g_{12}}) \cdots P(\rho_{g_{jm}}) \times \\ & P(\sigma^2_{b_1}) \cdots P(\sigma^2_{b_m}) P(\rho_{b_{12}}) \cdots P(\rho_{b_{jm}}) P(\sigma^2_{e_1}) \cdots P(\sigma^2_{e_m}) \end{aligned} \quad (1)$$

where $\mathbf{y} \mid \theta \sim N(\mu, \Sigma)$; the priors of the parameters are assumed to be independent and are assigned as follows.

I. $\mu_j \sim N(0, \sigma^2_\mu)$ is assigned to be a flat prior with very large variance;

II. $\sigma^2_{g_j} \sim \chi^{-2}(df, S_{g_j})$, we fix $df = 10$, and we assume that the genetic component in the model explains certain proportion of phenotypic variance by its genetic heritability, *i.e.*, $E(\sigma^2_{g_j}) = h^2_{g_j} * \text{var}(\mathbf{y}_j)$ where $h^2_{g_j}$ is the genetic heritability of a specific phenotypic trait in the $j^{th}$ environment; Then $E(\sigma^2_{g_j} \mid df, S_{g_j}) = \dfrac{df * S_{g_j}}{df - 2} = h^2_{g_j} * \text{var}(\mathbf{y}_j)$, thus

$S_{g_j} = \frac{h_{g_j}^2 * \text{var}(\mathbf{y}_j) * (df - 2)}{df}$. Similarly, we assume $\sigma_{b_j}^2 \sim \chi^{-2}(df, S_{b_j})$ and $\sigma_{e_j}^2 \sim \chi^{-2}(df, S_{e_j})$, set $df = 10$, then $S_{b_j} = \frac{h_{b_j}^2 * \text{var}(\mathbf{y}_j) * (df - 2)}{df}$ where $h_{b_j}^2$ is the background genetic heritability of a specific phenotypic trait in the $j^{th}$ environment; $S_{e_j} = \frac{(1 - h_{g_j}^2 - h_{b_j}^2) * \text{var}(\mathbf{y}_j) * (df - 2)}{df}$;

III. $\rho_{g_{jm}} \sim N(\mu_{g_{jm}}, \sigma^2)$ with condition $-1 \le \rho_{g_{jm}} \le 1$; Following the assumption that the expected value of genetic variance is a proportion of phenotypic variance, we assume $E(\sigma_{g_{jm}}) = h_{g_{jm}}^2 * \text{cov}(\mathbf{y}_j, \mathbf{y}_m)$, then $E(\rho_{g_{jm}} \sigma_{g_j} \sigma_{g_m}) = h_{g_{jm}}^2 * cor(\mathbf{y}_j, \mathbf{y}_m) * \sigma_{g_j} \sigma_{g_m}$, thus $\mu_{g_{jm}} = E(\rho_{g_{jm}}) = h_{g_{jm}}^2 * cor(\mathbf{y}_j, \mathbf{y}_m)$ where $h_{g_{jm}}^2$ is the genetic heritability of a specific phenotypic trait between the $j^{th}$ and $m^{th}$ environment; Likewise, we assume $\rho_{b_{jm}} \sim N(\mu_{b_{jm}}, \sigma^2)$ with condition $-1 \le \rho_{b_{jm}} \le 1$ and $\mu_{b_{jm}} = h_{b_{jm}}^2 * cor(\mathbf{y}_j, \mathbf{y}_m)$ where $h_{b_{jm}}^2$ is the background genetic heritability of a specific phenotypic trait across the $j^{th}$ and $m^{th}$ environment; the choice of $\sigma^2$ here is flexible, smaller $\sigma^2$ leads to a concentration on the expected value.

Due to the unknown form of the posterior conditional distribution of parameters, we adopt Random-walk Metropolis Hastings (RWMH) method to generate random samples from the full joint posterior distribution and then get Bayesian inference for estimated parameters. The basic idea of RWMH is to move the Markov chain with a certain acceptance probability by future candidate samples from a jumping distribution (*i.e.*, proposal distribution) that is conditional on the current state. The Markov chain proceeds in this random-walk way until the distribution of the current sample is close enough to the target distribution $P(\theta \mid \mathbf{y})$. There are two steps to execute a generic RWMH.

Step 1: Propose $\theta^* \sim g(\theta \mid \theta^{(t)})$, where $\theta^{(t)}$ is the current draw from $P(\theta \mid \mathbf{y})$;

Step 2: Accept $\theta^{(t+1)} = \theta^*$ if $u < \dfrac{P(\theta^* \mid \mathbf{y}) / g(\theta^* \mid \theta^{(t)})}{P(\theta^{(t)} \mid \mathbf{y}) / g(\theta^{(t)} \mid \theta^*)}$ where $u \sim Uniform(0,1)$;

Otherwise, go back to step 1.

In this study, we choose a symmetric proposal distribution, *i.e.*, $g(\cdot) = N(\theta^{(t)}, \sigma^2)$. Again, the choice of $\sigma^2$ is flexible, smaller $\sigma^2$ favors the proposal close to the current state. The model was implemented in software R with 100,000 iterations and the first 50,000 as burn-in for the interior spruce populations. The convergence of Markov chains was confirmed by visualizing the sampling paths.

**Table S1. Average mean squared error (standard deviation in parentheses) of genomic selection models for winter wheat DH population**. The results were collected over 50 replications of CV2 scheme; SE-GK , single-environment model with Gaussian kernel; ME-GK , multi-environment model with Gaussian kernel; ME-$WK_{MAF\text{-}Pvalue}$ (with $\beta = 12$), multi-environment model with WK by both MAF and p-value; GY, grain yield; SDS, SDS sedimentation value;, SKCSKW, kernel weight ; WHTPRO, wheat protein; E1/E2/E3=Year 2014/2015/2016. The significantly low values ($p<2.2\times10^{-16}$ by paired t-test) of each row are in boldface.

| **Trait** | **Environment** | **SE-GK** | **ME-GK** | **ME-$WK_{MAF\text{-}Pvalue}$** |
|---|---|---|---|---|
| GY | E1 | 0.71 (0.14) | 0.64 (0.13) | 0.62 (0.13) |
| | E2 | 0.74 (0.10) | 0.57 (0.08) | 0.57 (0.07) |
| | E3 | 0.87 (0.11) | 0.77 (0.10) | **0.38** (0.06) |
| SDS | E1 | 0.87 (0.13) | 0.64 (0.09) | **0.60** (0.09) |
| | E2 | 0.94 (0.14) | 0.49 (0.11) | 0.47 (0.11) |
| | E3 | 0.89 (0.10) | 0.45 (0.08) | **0.41** (0.07) |
| SKCSKW | E1 | 0.97 (0.14) | 0.85 (0.11) | 0.82 (0.10) |
| | E2 | 0.92 (0.16) | 0.74 (0.12) | **0.70** (0.11) |
| | E3 | 0.96 (0.12) | 0.70 (0.09) | **0.63** (0.08) |
| WHTPRO | E1 | 0.96 (0.15) | 0.85 (0.14) | **0.78** (0.12) |
| | E2 | 0.95 (0.11) | 0.94 (0.10) | 0.91 (0.11) |
| | E3 | 0.97 (0.13) | 0.86 (0.11) | **0.82** (0.11) |

**Table S2. Average mean squared error (standard deviation in parentheses) of genomic selection models for Interior spruce.** The results were collected over 50 replications of CV2 scheme; SE_GK , single-environment model with Gaussian kernel; ME_GK , multi-environment model with Gaussian kernel; $ME\_WK_{MAF\_Pvalue}$ (with $\beta = 200$), multi-environment model with WK by both MAF and p-value; HT, height; DBH, diameter at breast height; $WD_{res}$, resistance to drilling; $WD_{X\text{-}ray}$ , wood density in $kg/m^3$ using X-ray densitometry; E1/E2/E3 = PGTIS/Aleza Lake /Quesnel; PGTIS, Prince George Tree Improvement Station. The significantly low values ($p<2.2x10^{-16}$ by paired t-test) of each row are in boldface.

| Trait | Environment | SE-GK | ME-GK | $ME\text{-}WK_{MAF\text{-}Pvalue}$ |
|---|---|---|---|---|
| HT | E1 | 0.94 (0.15) | 0.98 (0.15) | **0.88** (0.15) |
| | E2 | 0.93 (0.12) | 0.90 (0.12) | **0.78** (0.11) |
| | E3 | 0.79 (0.11) | 0.81 (0.10) | **0.73** (0.10) |
| DBH | E1 | 0.99 (0.13) | 1.01 (0.13) | 0.99 (0.13) |
| | E2 | 0.98 (0.11) | 0.99 (0.11) | **0.85** (0.11) |
| | E3 | 0.94 (0.12) | 0.98 (0.12) | **0.90** (0.12) |
| $WD_{res}$ | E1 | 0.96 (0.13) | 0.96 (0.14) | **0.86** (0.13) |
| | E2 | 0.90 (0.10) | 0.91 (0.10) | **0.80** (0.10) |
| | E3 | 0.94 (0.12) | 0.96 (0.12) | **0.88** (0.12) |
| $WD_{X\text{-}ray}$ | E1 | 0.98 (0.16) | 0.97 (0.17) | 0.96 (0.17) |
| | E2 | 0.94 (0.11) | 0.94 (0.11) | **0.89** (0.10) |
| | E3 | 0.94 (0.12) | 0.94 (0.11) | **0.90** (0.11) |

**Table S3. The estimated variance component proportions from genomic selection models for winter wheat DH population.** SE-GK , single-environment model with Gaussian kernel; ME-GK , multi-environment model with Gaussian kernel; ME-WK$_{\text{MAF-Pvalue}}$ (with $\beta = 12$), multi-environment model with WK by both MAF and p-value; GY, grain yield; SDS, SDS sedimentation value; SKCSKW, kernel weight; WHTPRO, wheat protein; E1/E2/E3=Year 2014/2015/2016; $\sigma_g^2$, main genetic variance; $\sigma_b^2$, background genetic variance; $\sigma_e^2$, residual variance.

| | | GY | | | SDS | | |
|---|---|---|---|---|---|---|---|
| | | **SE-GK** | **ME-GK** | **ME-WK$_{\text{MAF-Pvalue}}$** | **SE-GK** | **ME-GK** | **ME-WK$_{\text{MAF-Pvalue}}$** |
| **E1** | $\sigma_g^2$ | 0.72 | 0.75 | 0.84 | 0.58 | 0.73 | 0.80 |
| | $\sigma_b^2$ | 0.11 | 0.11 | 0.07 | 0.17 | 0.12 | 0.09 |
| | $\sigma_e^2$ | 0.17 | 0.14 | 0.09 | 0.25 | 0.15 | 0.11 |
| **E2** | $\sigma_g^2$ | 0.73 | 0.79 | 0.86 | 0.48 | 0.71 | 0.83 |
| | $\sigma_b^2$ | 0.11 | 0.11 | 0.07 | 0.21 | 0.18 | 0.08 |
| | $\sigma_e^2$ | 0.16 | 0.10 | 0.07 | 0.31 | 0.11 | 0.09 |
| **E3** | $\sigma_g^2$ | 0.74 | 0.77 | 0.95 | 0.61 | 0.78 | 0.88 |
| | $\sigma_b^2$ | 0.11 | 0.10 | 0.02 | 0.17 | 0.14 | 0.06 |
| | $\sigma_e^2$ | 0.15 | 0.13 | 0.03 | 0.22 | 0.08 | 0.06 |
| | | **SKCSKW** | | | **WHTPRO** | | |
| **E1** | $\sigma_g^2$ | 0.37 | 0.42 | 0.87 | 0.41 | 0.47 | 0.78 |
| | $\sigma_b^2$ | 0.24 | 0.28 | 0.05 | 0.23 | 0.28 | 0.10 |
| | $\sigma_e^2$ | 0.39 | 0.30 | 0.08 | 0.36 | 0.25 | 0.12 |
| **E2** | $\sigma_g^2$ | 0.35 | 0.36 | 0.83 | 0.41 | 0.48 | 0.79 |
| | $\sigma_b^2$ | 0.25 | 0.40 | 0.07 | 0.24 | 0.21 | 0.10 |
| | $\sigma_e^2$ | 0.40 | 0.24 | 0.10 | 0.35 | 0.31 | 0.11 |
| **E3** | $\sigma_g^2$ | 0.38 | 0.41 | 0.89 | 0.25 | 0.30 | 0.66 |
| | $\sigma_b^2$ | 0.24 | 0.40 | 0.05 | 0.28 | 0.37 | 0.13 |
| | $\sigma_e^2$ | 0.38 | 0.19 | 0.06 | 0.47 | 0.33 | 0.21 |

**Table S4. The estimated variance component proportions from genomic selection models for Interior spruce.** SE_GK , single-environment model with Gaussian kernel; ME_GK , multi-environment model with Gaussian kernel; $ME\_WK_{MAF\_Pvalue}$ (with $\beta = 200$), multi-environment model with WK by both MAF and p-value; HT, height; DBH, diameter at breast height; $WD_{res}$, resistance to drilling; $WD_{X\text{-}ray}$ , wood density in $kg/m^3$ using X-ray densitometry; E1/E2/E3 = PGTIS/Aleza Lake /Quesnel; PGTIS, Prince George Tree Improvement Station; $\sigma_g^2$**,** main genetic variance; $\sigma_b^2$**,** background genetic variance; $\sigma_e^2$**,** residual variance.

| | | HT | | | DBH | | |
|---|---|---|---|---|---|---|---|
| | | **SE-GK** | **ME-GK** | **$ME\text{-}WK_{MAF\text{-}Pvalue}$** | **SE-GK** | **ME-GK** | **$ME\text{-}WK_{MAF\text{-}Pvalue}$** |
| **E1** | $\sigma_g^2$ | 0.40 | 0.31 | 0.49 | 0.35 | 0.25 | 0.35 |
| | $\sigma_b^2$ | 0.28 | 0.35 | 0.22 | 0.27 | 0.27 | 0.26 |
| | $\sigma_e^2$ | 0.32 | 0.34 | 0.29 | 0.38 | 0.48 | 0.39 |
| **E2** | $\sigma_g^2$ | 0.30 | 0.23 | 0.60 | 0.37 | 0.25 | 0.47 |
| | $\sigma_b^2$ | 0.32 | 0.29 | 0.15 | 0.23 | 0.24 | 0.20 |
| | $\sigma_e^2$ | 0.38 | 0.48 | 0.25 | 0.40 | 0.46 | 0.33 |
| **E3** | $\sigma_g^2$ | 0.51 | 0.34 | 0.57 | 0.29 | 0.29 | 0.36 |
| | $\sigma_b^2$ | 0.24 | 0.51 | 0.29 | 0.31 | 0.34 | 0.32 |
| | $\sigma_e^2$ | 0.25 | 0.15 | 0.14 | 0.40 | 0.37 | 0.32 |
| | | | **$WD_{res}$** | | | **$WD_{X\text{-}ray}$** | |
| **E1** | $\sigma_g^2$ | 0.28 | 0.31 | 0.55 | 0.27 | 0.26 | 0.28 |
| | $\sigma_b^2$ | 0.34 | 0.34 | 0.20 | 0.27 | 0.31 | 0.33 |
| | $\sigma_e^2$ | 0.38 | 0.35 | 0.25 | 0.46 | 0.43 | 0.39 |
| **E2** | $\sigma_g^2$ | 0.28 | 0.22 | 0.54 | 0.28 | 0.26 | 0.48 |
| | $\sigma_b^2$ | 0.40 | 0.39 | 0.19 | 0.30 | 0.30 | 0.21 |
| | $\sigma_e^2$ | 0.32 | 0.39 | 0.27 | 0.42 | 0.44 | 0.31 |
| **E3** | $\sigma_g^2$ | 0.35 | 0.27 | 0.38 | 0.26 | 0.25 | 0.34 |
| | $\sigma_b^2$ | 0.27 | 0.27 | 0.22 | 0.33 | 0.32 | 0.27 |
| | $\sigma_e^2$ | 0.38 | 0.46 | 0.40 | 0.41 | 0.43 | 0.39 |

**Fig. S1. The relationship between minor allele frequency and the weight from minor allele frequency by different values of $\beta$ for winter wheat DH population and site PGTIS of Interior spruce.** PGTIS, Prince George Tree Improvement Station.

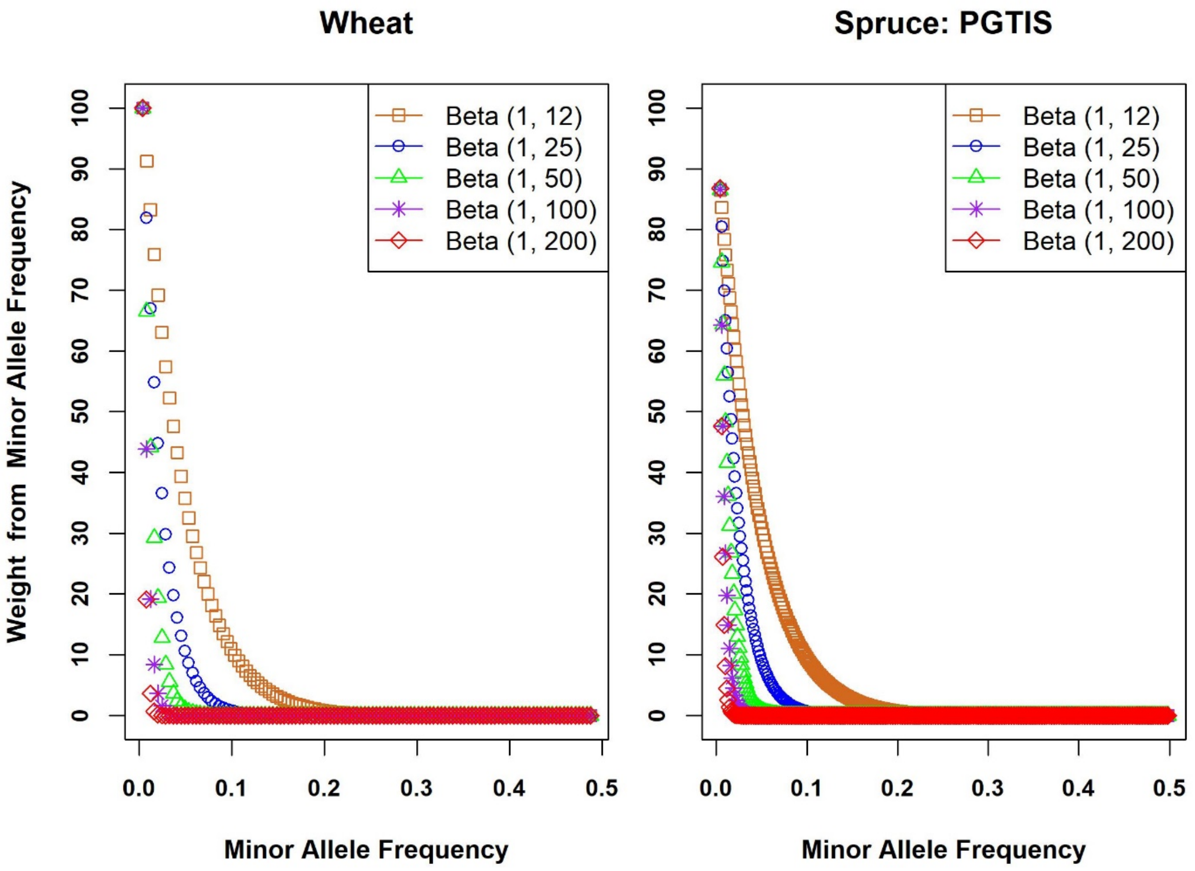

**Fig. S2. Average Pearson's correlation coefficients over 50 replications of CV2 scheme from multi-environment model with weighted kernel by both minor allele frequency and p-value ($ME\_WK_{MAF\_Pvalue}$) and from different values of $\beta$ for winter wheat DH population.** GY, grain yield; SDS, SDS sedimentation value; SKCSKW, kernel weight; WHTPRO, wheat protein.

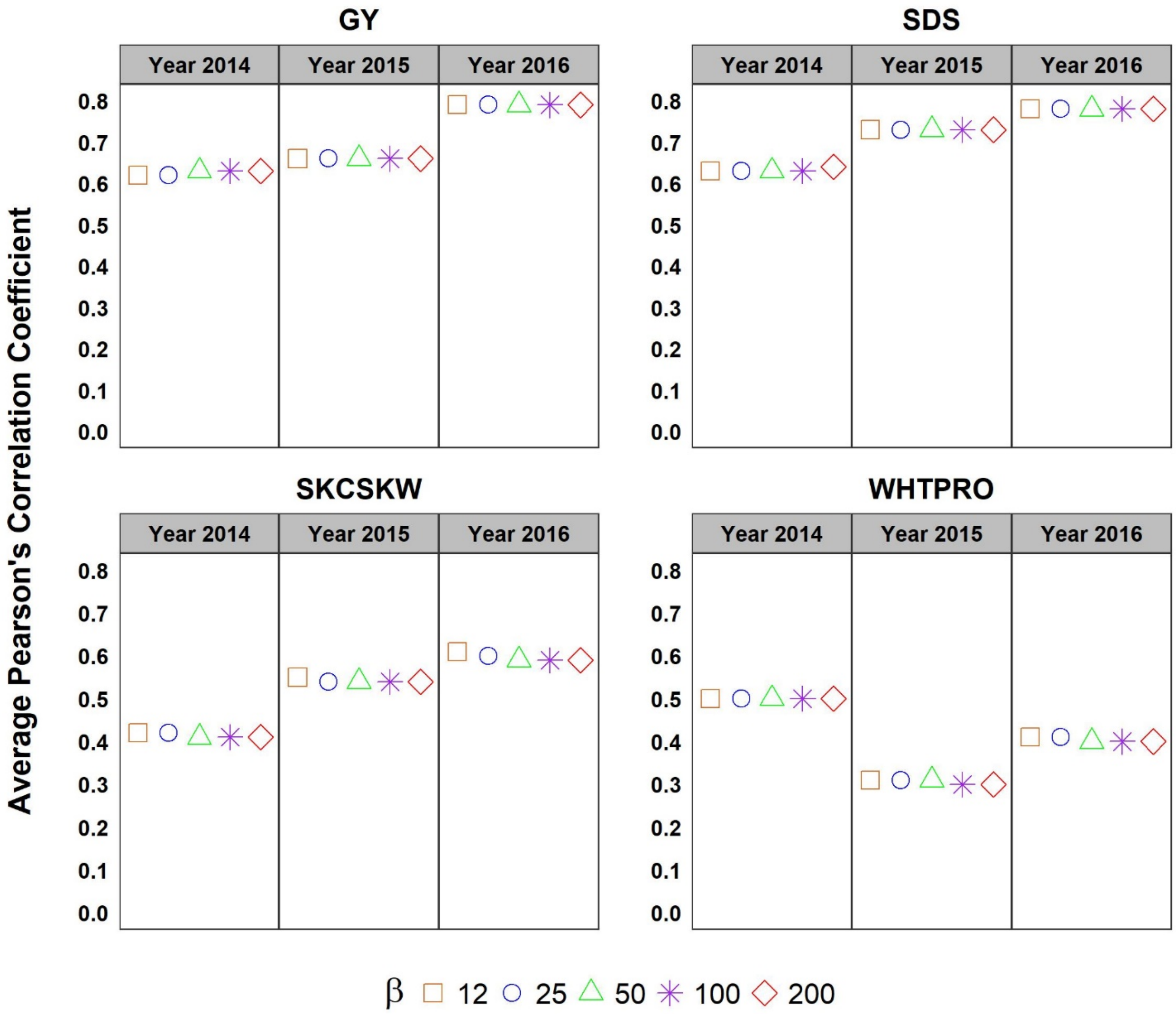

**Fig. S3. Average Pearson's correlation coefficients over 50 replications of CV2 scheme from multi-environment model with weighted kernel by both minor allele frequency and p-value ($ME\_WK_{MAF\_Pvalue}$) and from different values of $\beta$ for Interior spruce.** HT, height; DBH, diameter at breast height; $WD_{res}$, resistance to drilling; $WD_{X\text{-}ray}$, wood density in kg/m$^3$ using X-ray densitometry; PGTIS, Prince George Tree Improvement Station.

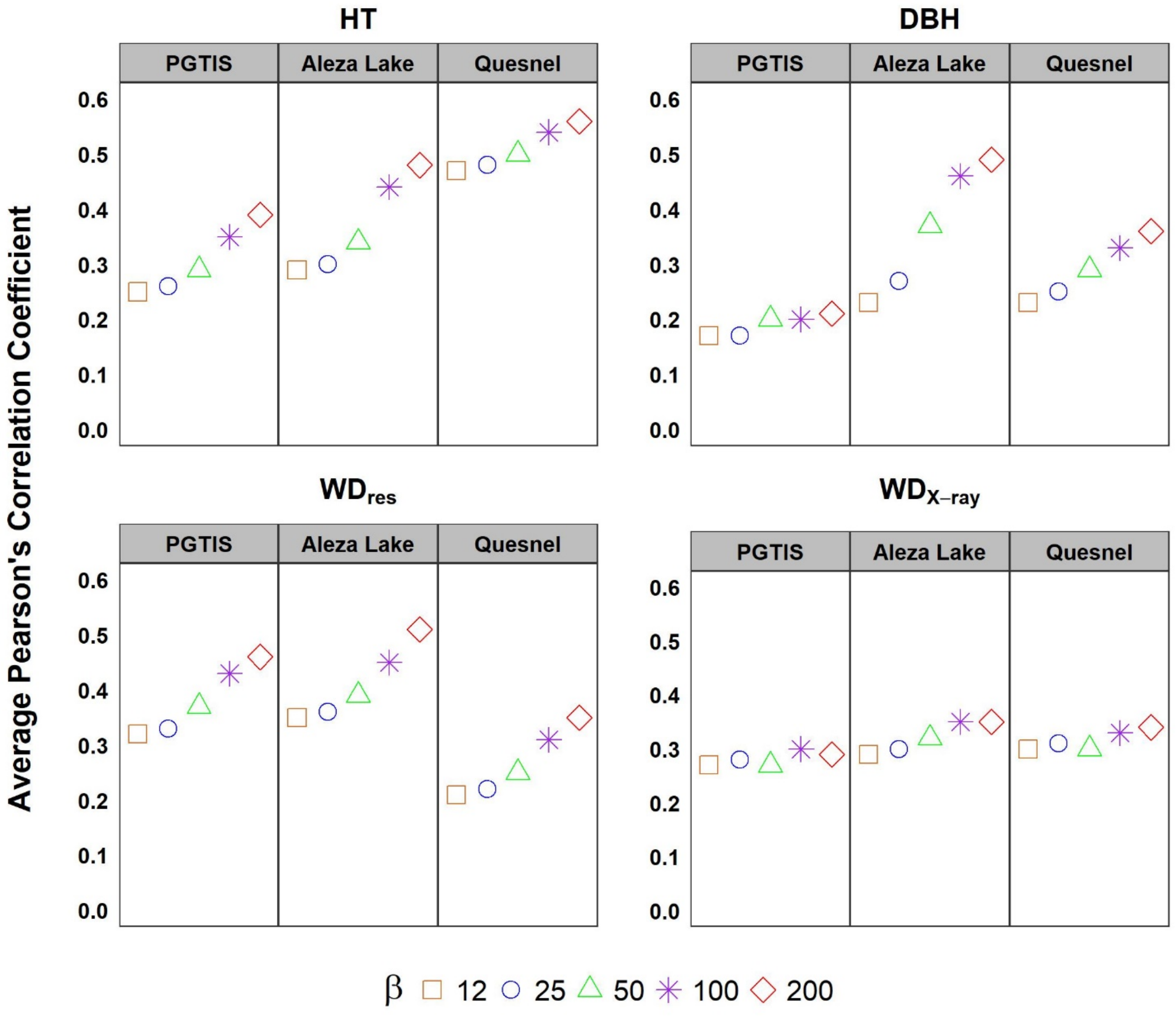
